# Contrast Sensitivity Function in Deep Networks

**DOI:** 10.1101/2023.01.06.523034

**Authors:** Arash Akbarinia, Yaniv Morgenstern, Karl R. Gegenfurtner

## Abstract

The contrast sensitivity function (CSF) is a fundamental signature of the visual system that has been measured extensively in several species. It is defined by the visibility threshold for sinusoidal gratings at all spatial frequencies. Here, we investigated the CSF in deep neural networks using the same 2AFC contrast detection paradigm as in human psychophysics. We examined 240 networks pretrained on several tasks. To obtain their corresponding CSFs, we trained a linear classifier on top of the extracted features from frozen pretrained networks. The linear classifier is exclusively trained on a contrast discrimination task with natural images. It has to find which of the two input images has higher contrast. The network’s CSF is measured by detecting which one of two images contains a sinusoidal grating of varying orientation and spatial frequency. Our results demonstrate characteristics of the human CSF are manifested in deep networks both in the luminance channel (a band-limited inverted U-shaped function) and in the chromatic channels (two low-pass functions of similar properties). The exact shape of the networks’ CSF appears to be task-dependent. The human CSF is better captured by networks trained on low-level visual tasks such as image-denoising or autoencoding. However, human-like CSF also emerges in mid- and high-level tasks such as edge detection and object recognition. Our analysis shows that human-like CSF appears in all architectures but at different depths of processing, some at early layers, while others in intermediate and final layers. Overall, these results suggest that (i) deep networks model the human CSF faithfully, making them suitable candidates for applications of image quality and compression, (ii) efficient/purposeful processing of the natural world drives the CSF shape, and (iii) visual representation from all levels of visual hierarchy contribute to the tuning curve of the CSF, in turn implying a function which we intuitively think of as modulated by low-level visual features may arise as a consequence of pooling from a larger set of neurons at all levels of the visual system.

## 1. Introduction

Normalising the input signals relative to the context plays a major role during the transformation from sensation to perception (Carandini and Heeger, 2012). In human visual perception, for example, our ability to perceive a visual scene is determined by the relative contrast between the constituent details. Contrast is broadly defined as the relative difference between the foreground and background (Peli, 1990; Pelli and Bex, 2013). Our ability to detect contrast is a primary step in biological image processing that eventually leads to functional behaviours, like object detection, scene recognition, and semantic segmentation.

At a behavioural or systems level, one of the most fundamental signatures of biological visual processing is the luminance contrast sensitivity function (CSF) (Schade, 1956; Campbell and Robson, 1968). The CSF is based on human contrast thresholds over a wide range of spatial frequencies. Their results show that the luminance-CSF is a band-limited inverted U-shaped function with lower sensitivity at high and low frequencies. Since then, numerous other studies have consistently reproduced the human CSF, e.g., (Peli et al., 1996; Hashemi et al., 2012; Kim and Kim, 2010), for review see (Graham, 1989). Another large body of work has examined the CSF across many different species of animals obtaining the typical inverted U-shape function with some shifts in the peak of spatial frequency according to the species’ visual system, e.g., (Harmening et al., 2009; Bisti and Maffei, 1974; De Valois et al., 1974; Northmore and Dvorak, 1979; Hirsch, 1982; Reymond and Wolfe, 1981; Hodos et al., 2002).

The CSF has been applied and extended in many ways. In clinical settings, the CSF is used to test visual acuity, assess visual dysfunction, and monitor ophthalmological treatments (Owsley, 2003). In computer vision, the CSF has also been proposed as a tool to assess image quality (Wang et al., 2004; Kim and Lee, 2017). Regarding chromatic contrast sensitivity, chromatic-CSFs for both red-green and yellow-blue channels are low-pass functions with no attenuation even in the lowest producible spatial frequency (Kelly, 1983; Mullen, 1985).

Campbell and Robson (1968) suggested that CSF arises because of the presence of multiple channels in the visual system, each selective to a different band of spatial frequencies. Others have explained the low-frequency fall-off solely by neural factors, such as the extent of lateral inhibition (Wandell, 1995; Cornsweet, 1970), or specific properties of low-frequency channels (Graham, 1972), e.g., their relative insensitivity to low spatial frequency channels. Barten (1999) proposed that a low-frequency fall-off is an approach the visual system learns to efficiently use the signals’ dynamic range, given the system’s physical capacity. The high-frequency drop is explained based on a combination of physical constraints leading to optical degradation (e.g., ocular aberrations) and receptor cell spacing in the retina (Campbell and Green, 1965; Cornsweet, 1970).

These explanations of CSF suggest early visual processes are the basis of distinct visibility thresholds at different spatial frequencies; indeed, our ability to process contrast begins with single neurons in the retina and lateral geniculate nucleus excited most by light in the centre coinciding with dark in the surroundings or vice versa (Kuffler, 1953; Hubel and Wiesel, 1961). In the early visual cortex, neurons become sensitive to patterns of light that specify changes in orientation, spatial frequency, motion, and colour (Hubel and Wiesel, 2004) with highly localised responses strongly modulated by contrast; such single neurons are thought to underlie the human CSF and to efficiently code our environments (Atick et al., 1992; Atick and Redlich, 1992; Atick, 1992; Li et al., 2022; Gomez-Villa et al., 2020). Although there is no doubt about the advantages of efficient visual processing (Geisler, 2008; Olshausen and Field, 1996), it is unclear how other important evolutionary behaviours (such as high-level visual tasks like object recognition) have affected the emergence of CSF in biological systems.

Deep neural networks (DNNs) trained on the ImageNet dataset (Deng et al., 2009) to classify images into one thousand categories exhibit several characteristics of object processing in the ventral stream, e.g., (Bashivan et al., 2019; Cadieu et al., 2014; Cichy et al., 2016; Eickenberg et al., 2017; Akbarinia and Gil-Rodŕıguez, 2021; Flachot et al., 2022; Storrs et al., 2021; de Vries et al., 2022). Similar to biological neurons, the artificial kernels at the early level of these networks have Gabor-like receptive fields tuned to orientation and spatial frequency; the kernels’ receptive field becomes larger in deeper layers, representing progressively more complex visual patterns (e.g., shapes, and objects), and predicts human brain responses from fMRI in area IT, e.g., (Yamins et al., 2014; Khaligh-Razavi and Kriegeskorte, 2014), for review see (Lindsay, 2020). We take advantage of this striking resemblance between the properties of artificial neurons in these networks and biology to examine how filtering responses at different layers can serve as a proxy of systems-level responses in humans towards contrast (See Section 3 Object Recognition).

Object recognition is only one of many critical behavioural tasks for the survival of biological organisms; thus, we also investigate the CSF of deep networks trained for other behavioural tasks (e.g., noise removal, edge detection, scene segmentation, etc.) by studying the Taskonomy networks (an identical architecture trained on the same set of images for different visual tasks). The comparison across this set of networks, whose difference originates from the optimisation of distinct loss functions according to task demand, allows us to assess how task-dependent features shape the CSF of a visual system. Complementary to the current explanations of human CSF that emerge due to low-level features for efficient coding (Barlow et al., 1961), our results suggest that human-like CSF appears at all hierarchical levels of visual representation. In addition, we show that networks trained on ecologically valid tasks produce more human-like CSF than more artificial tasks.

## 2. Method

The CSF is typically measured with a 2AFC (two-alternative forced choice) or 2IFC (two-interval forced choice) psychophysical paradigm which presents humans with two stimuli. One stimulus contains a sinusoidal grating while the other is a uniform patch of zero contrast. Participants report the image with the visible grating (left or right in Figure 1-a). The grating contrast is adjusted by a staircase procedure and observer responses are then used to estimate a participant’s performance at 75% correct. Sensitivity is defined as the inverse of contrast.

**Figure 1:**
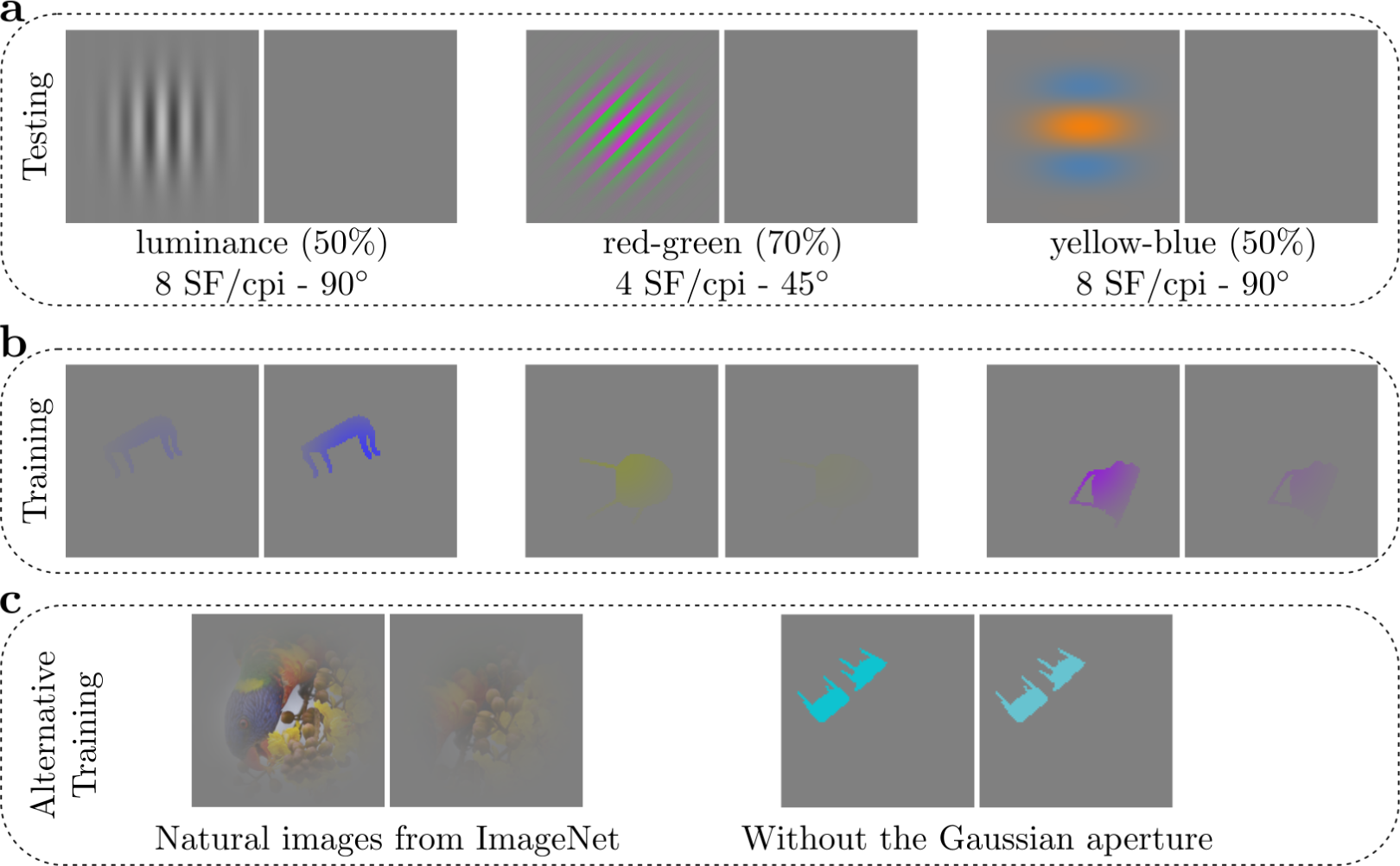
Example stimuli used in this study. **a**: sinusoidal gratings to measure CSF; network report which image contains grating. **b**: natural objects to train contrast discriminator linear classifier; network report which image has higher contrast. **c**: other explored options to train the linear classifier.

For a neural network trained on a task like object recognition measuring the CSF for this 2AFC task is impossible, as the neural network was specifically trained for another task. We can, however, use a linear classifier (Alain and Bengio, 2017) to evaluate how well a neural network’s already evolved features can learn to do the 2AFC contrast detection task. That is to say, the framework to evaluate the CSF in deep networks consists of two steps. First, a network is trained on an arbitrary visual task (e.g., object recognition). We refer to such a network as a **pretrained network**. Second, features extracted from the pretrained network are input to a linear classifier trained for the contrast discrimination 2AFC task. We refer to the trained linear classifier as a **contrast-discriminator**. Thus, the aim is to compute the CSF for the features learned by an artificial neural network performing a visual task, i.e., the pretrained network. The contrast-discriminator is a way to assess the CSF as we cannot directly ask the pretrained network which image has higher contrast. Figure 2-a illustrates the schematic of this procedure; features are extracted from a frozen pretrained network, and only the weights of the linear classifier are optimised during the contrast-discrimination task.

**Figure 2:**
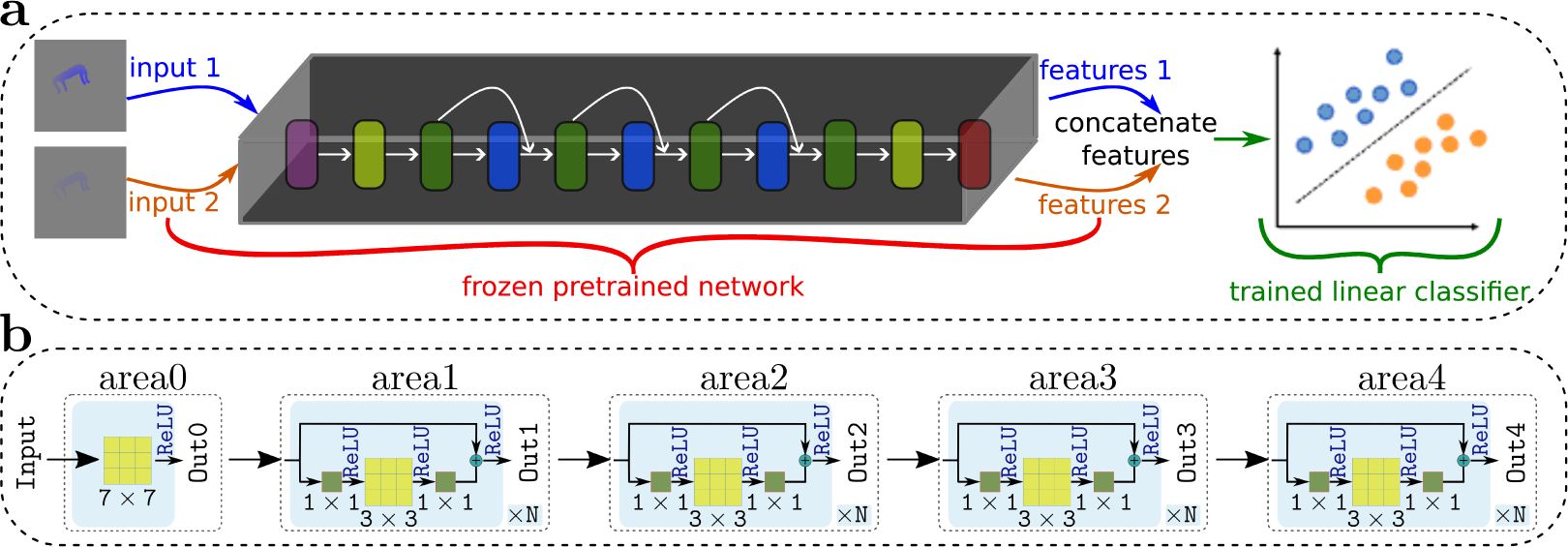
**a**: The schematic flowchart of our framework. Features can be extracted from any arbitrary pretrained network at any intermediate layer. As an example, using the ResNet architecture in **b**, the frozen pretrained network can take on any of the *area0–4* building blocks. In this study, we explored more than one hundred networks performing several different tasks. **b**: The ResNet architecture. The building block of *area1-4* is the same only the number of times they repeat (*N_i_*) can differ. In ResNet50, *N*_1_ = 3, *N*_2_ = 4, *N*_3_ = 6, and *N*_4_ = 3.

### 2.1. Pretrained networks

We studied 70 different architectures. The majority of those are convolutional neural networks (CNNs) (Krizhevsky et al., 2012). A few are vision transformers (ViTs) (Vaswani et al., 2017). CNNs process the input image by convolution kernels that slide along input features followed by non-linear rectification and downsampling operations. ViTs, on the other hand, divide the input into a sequence of small patches (e.g., 16 *×* 16) and computes relationships among these image regions. All architectures except AlexNet and VGG incorporate *skip connections* similar to the biological brain in which intermediary layers are skipped across or within areas (Thomson, 2010).

To investigate the CSF of networks’ internal representation, we extracted features by a forward pass of an image (i.e., the pretrained weights are frozen without updating them by a backward error propagation). To examine how the CSF develops at different layers, we extracted features at different levels of visual hierarchy. Figure 2-a depicts the feature extraction process from the last layer of a network, which can also be applied to intermediate layers.

We examined the *ResNet* architecture (He et al., 2016) in greater detail for two reasons: (i) the model’s depth is parametrically adjustable according to the number of residual blocks, and (ii) the same architecture has been employed as the backbone for several other tasks, e.g., semantic segmentation (Chen et al., 2018), edge detection, denoising (Zamir et al., 2018), or vision-language models (Radford et al., 2021). Given the availability of many pretrained ResNet models (i.e., identical architecture with different weights optimised for specific tasks), we can examine how the network’s task modulates its CSF.

The ResNet architecture consists of five *areas* (see Figure 2-b). The initial input area (*area0*) is fixed by a convolutional layer of kernel size 7 *×* 7 and a max pooling layer over a region of 3 *×* 3. The other four areas encompass several residual blocks whose skip connections are modelled by summing the input to the output. Within each area, the spatial resolution remains intact. Across areas, the spatial resolution of the signal is reduced by a factor of two as a result of step size in the convolution operation.

### 2.2. Contrast-discriminator linear classifier

We trained the contrast discriminator linear classifier exclusively with images of real-world objects randomly placed/coloured on a uniform background of random luminance (see Figure 1-b). During the training, the images are presented in the *uint8* precision making with the maximum possible contrast 0.392% (^1^). Images are multiplied by a Gaussian aperture to smooth the transition from foreground to background. The training stimuli consist of two images of random contrast in the range of [0, 1]. In half of the samples, the image contrast was modulated equally in all three RGB channels; in the other half, we modulated only the contrast of one channel (R, G, B). The rationale is to mimic real-world scenarios where sometimes the contrast of all photoreceptors is simultaneously affected, and sometimes only one photoreceptor. Nevertheless, we also explored two other versions in which (i) always all RGB channels were equally modified or (ii) always only one channel was modified. The pattern of results remains very similar.

In another set of experiments, we trained the linear classifier with natural images from the *ImageNet* dataset (Deng et al., 2009), a large visual database spanning over 1000 object categories. We also explored the option of removing the Gaussian aperture (Figure 1-c). The patterns of results remained almost identical in both cases. Overall, the linear classifier, as we will discuss more in-depth later, does not impact the obtained results. It is merely a tool to interpret the features of a pretrained network.

Figure 2-a depicts the schematic of our framework. We input the pretrained network with one image and extract its features from a specific layer (this procedure is also known as readouts). The extracted features are vectorised irrespective of the spatial resolution of the feature map. We repeat the same procedure for the second image. We concatenate in random order the extracted features for the low- and high-contrast images into a single vector, which serves as input to a linear classifier. The network performs a 2AFC task to identify which image has higher contrast. We trained the linear classifiers in *PyTorch* with identical settings (e.g., the same weight initialisation and an identical random seed) across all training instances^1^. Input images are colour (RGB) images of size 224 *×* 224 augmented with random horizontal flips and cropping. The linear classifier is trained on a single GPU with a batch size of 32 for ten epochs of 15K samples (i.e., 150K iterations). We used the stochastic gradient descent (SGD) optimiser and updated the weights of the linear classifier by computing the binary cross-entropy loss. Within this process, the weights of the pretrained network are frozen(i.e., not updated).

### 2.3. Measuring the CSF

We opted for the contrast detection task similar to the ModelFest dataset (Carney et al., 1999), aiming to measure the networks’ contrast sensitivity function (CSF) as closely as possible to human psychophysics. Figure 1-a presents example stimuli from three trials. Each trial consisted of two intervals, where one interval showed an image with a non-zero contrast modulated sinusoidal gratings, and the other showed an image of the uniform grey background of 0% contrast. During the testing, the images are presented in *floating* precision to assess the full sensitivity of a network. The image sizes were 224 *×* 224 pixels. The network’s task was to select the interval with the grating. The sinusoidal grating contrast was adjusted with a staircase procedure until the network reached 75% correct - the same threshold typically assumed in human experiments.

We compute the network’s accuracy across sixteen conditions: four orientations (0, 45, 90 and 135*^◦^*), two phases (0 and 180) and two image orders (first or second corresponding to left or right images in Figure 1-a). An image of size 224 *×* 224 allows for exact spatial frequencies of [1, 2, 4, 7, 8, 14, 16, 28, 32, 56, 112] cycles per image (cpi). We chose these exact settings to avoid any artefacts in test images. For instance, other spatial frequencies cannot be exactly produced (if they are not divisible to 224). Similarly, other orientations need interpolation across different pixels and are therefore avoided. We generated the **luminance-gratings** by modulating all RGB channels with the same contrast level, which is numerically identical to modulating only the luminance channel of a colour-opponent space such as DKL.

**Chromatic-gratings**: The networks’ input is in the RGB colour space (given that the pretrained networks were trained with RGB images). To avoid artefacts caused by converting to RGB, we relied on the YP*_b_*P*_r_* colour space instead of a biologically plausible colour space like DKL, where contrast reduction in one channel of DKL causes different levels of contrast modulation in the RGB channels (See, for example, Figure A.1 in appendices). The contrast modulation we performed in the YP*_b_*P*_r_* results in sinusoidal gratings of identical waves with different phases for the R- and G- and B-channels (Mullen, 1985).

### 2.4. Comparison to human CSF

**Assumption**: In an image of size 224 *×* 224, the maximum spatial frequency we can generate is 112 cpi and the minimum 1. The human fovea spans over two visual degrees with a 60 cycles/deg resolution. To compare the results of networks to human data, we assumed the networks’ field of view is equivalent to the entire fovea of the human visual system (2*^◦^*). Therefore, we assumed one cycle/image is equivalent to half a cycle/deg.

We quantitatively compare the networks’ luminance-CSF to the human data from (Campbell and Robson, 1968) and the chromatic-CSF to the data from (Mullen, 1985) using the Pearson correlation coefficient. We used a model of a double exponential function to obtain the human luminance-CSF (Uhlrich et al., 1981) that has also been reported a good fit to the CSF of other animals (Harmening et al., 2009). Correlation measures naturally ignore the amplitude of the CSF and evaluate only the shape of the function.

We also actively analyse networks CSF graphically to prevent the effect of outliers and influential data points on the correlation coefficient (Anscombe, 1973). To do this, the network’s CSF is normalised (divided) by the maximum sensitivity of the network across three channels (luminance, red-green, yellow-blue), which only brings the maximum amplitude to one while keeping the ratio across channels intact.

## 3. Object recognition

ImageNet pretrained networks have been extensively compared to human vision (Bashivan et al., 2019; Cadieu et al., 2014; Cichy et al., 2016; Eickenberg et al., 2017; Khaligh-Razavi and Kriegeskorte, 2014; Storrs et al., 2021; Yamins et al., 2014; Zeman et al., 2020). Thus, we first explored the CSF of several networks (available from the PyTorch framework^2^) that have been trained on the ImageNet (Deng et al., 2009) dataset to perform object classification across 1000 categories.

### 3.1. Classification layer

The last layer of an ImageNet pretrained network (also known as the classification layer) is of identical size for all architectures (i.e., a vector of length 1000), which facilitates a straightforward comparison across networks’ classification feature space. We extracted features from the classification layer of 126 networks (from 63 architectures) and trained a contrast-discriminator linear classifier on top of each (see Figure 2). Some networks’ CSF is highly non-systematic (e.g., ConvNeXt; Figure 3-a). Some networks capture the human chromatic-CSF very well, but their luminance-CSF does not resemble the human CSF (e.g., ViT-L32, *r_rg_* = 0.96 and *r_yb_*= 0.98). Another group only obtains a human-like luminance-CSF (e.g., RegNet, *r* = 0.92 in luminance channel). A few other networks resemble both chromatic- and luminance-CSF of humans (e.g., ResNet50, *r_µ_* = 0.84 across three channels). The average CSF (i.e., the CSF obtained by averaging the sensitivity of all individual instances) resembles the human CSF both qualitatively (inverted-U shape function in the luminance channel, and two low-pass filters of similar properties in chromatic channels) and quantitatively (*r_µ_* = 0.69 across three channels). The entire distribution of correlations to the human CSF for individual networks and one instance of each of the 63 ImageNet architectures are reported in Figure 3-c–d, respectively. Overall, there is a large discrepancy between the CSF across networks with varied architectures.

**Figure 3:**
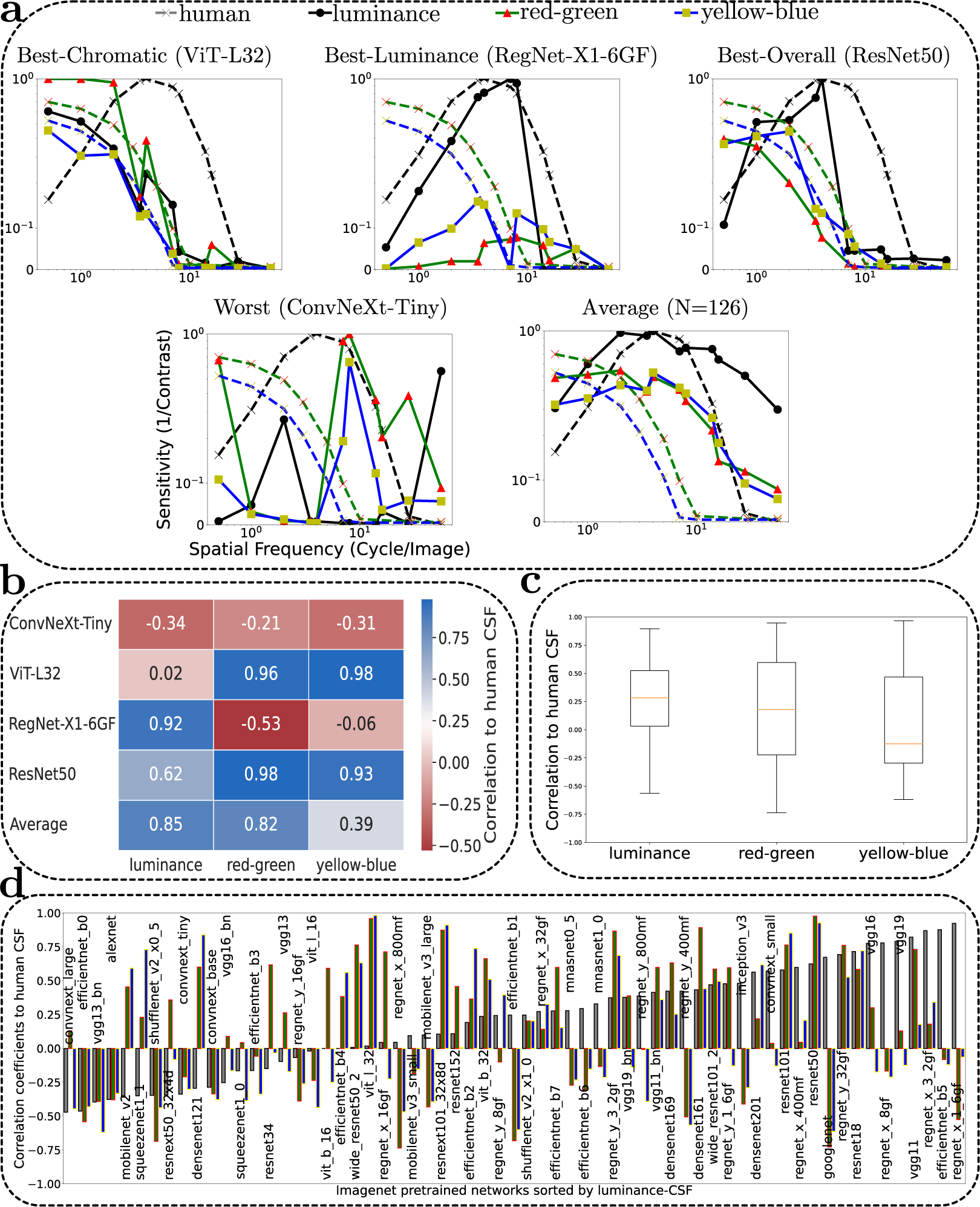
**a**: The CSF of four ImageNet pretrained networks by reading out features from the final layer (i.e, fc, object classification layer). **b**: The correlation to the human CSF. **c**: The distribution of correlation to the human CSF for 126 networks (from 63 architectures). The box extends from the first to third quartile values of the data, with an orange line at the median. **d**: The correlation of 63 ImageNet pretrained networks to human CSF sorted by the correlation to the luminance CSF. Grey bars indicate the luminance-CSF, and green and blue bars are the chromatic-CSF (red-green and yellow-blue, respectively).

We examined whether there is a relationship between the degree of similarity to the human CSF and the networks’ accuracy in the pretrained tasks (i.e., object recognition), or the networks’ performance when image contrast is reduced (Geirhos et al., 2018; Akbarinia and Gil-Rodriguez, 2020), but we found no correlation (luminance: *r* = 0*, p* = 0.97; red-green: *r* = 0.18*, p* = 0.3; yellow-blue: *r* = 0.04*, p* = 0.8). Neither the depth of the network nor its number of units explains the degree of similarity to the human CSF. Furthermore, within the same family of architecture, some networks might obtain human-like CSF while others do not. For instance, VGG11/16/19 exhibit human-like luminance-CSF but not VGG13, similarly the CSF of ResNet18/50/101 in all three channels corresponds very well to the human CSF, but this is not the case for ResNet34. Two hypotheses can be behind these discrepancies. First, human-like CSF in object recognition networks only emerges as a result of some particular evolution of features during the training procedure. Second, human-like CSF might appear in different layers, not necessarily in the final layer. While these two hypotheses are not mutually exclusive, we will show evidence for the latter hypothesis in the next section.

It is important to emphasise that we did not introduce any constraints on these networks to obtain human-like CSF. It emerges as a consequence of training the contrast-discriminator with features that are required for successful object recognition in natural images. The average results across all instances suggest similarities to human CSF: (i) the networks’ luminance-CSF is band-pass with a peak of its sensitivity matching the peak of human CSF, (ii) the networks’ chromatic-CSF is low-pass whose sensitivity steadily attenuates as a function of spatial frequency, and (iii) the amplitude of the networks’ luminance-CSF is larger than its chromatic-CSF. Nevertheless, there are also differences between human and networks’ CSF: (i) the human luminance-CSF drops sharply to zero in high-SFs, while the average CSF of networks never drops to zero in the highest spatial frequency, (ii) the human chromatic-CSF drops sharply to zero in high-SFs, while the average CSF of networks never drops to zero in the highest spatial frequency, (ii) the human chromatic-CSF drops sharply to zero in mid-SFs, and (iii) the amplitude ratio between luminance- and chromatic-CSF is larger in humans. Furthermore, the absolute sensitivity (1/threshold) in networks is considerably larger than in humans, which is in the range of 100 *−* 200 depending on the luminance or chromatic gratings. The maximum amplitude for networks is 20*K* (luminance), 5.2*K* (red-green) and 8.8*K* (yellow-blue), and the median sensitivity is 582 (luminance), 275 (red-green) and 261 (yellow-blue). The lowest value of the random contrast in our training is 0.4%. If this factor limited the network’s feature space, the maximum sensitivity would be about 255. However, it seems networks become hyper-sensitive at test time (with images of *floating* points) perhaps by pooling across pixels.

### 3.2. Internal representation

Next, we took a subset of these networks that obtained the lowest and highest correlations to the human CSF, covering both CNN and ViT architecture (i.e., ResNet50, VGG16, ViT-B32, RegNet, ConvNeXt, and ViT-L32) and analysed their internal representation to reveal which layers have features that produce the shape of CSF. Figure 4 (and B.1 in appendices) report the CSF of these networks at five intermediate depths and the last classification layer (*fc*). We matched the intermediate layers across networks by selecting these layers at similar depths (however, as the networks have different architectures, there is no exact match). A human-like CSF emerges in all networks, with ResNet50 best capturing the CSF across its areas. In addition to a high correlation to the human CSF (*r ≥* 0.89 in all three channels), qualitatively, the ResNet50’s CSF shows a clear preference for some subbands (the CSF shape is less noisy) even in areas where the CSF is unlike humans (e.g., *area0*). The peak of its luminance-CSF (in *area3–4*) lies precisely on the peak of human luminance-CSF and its sensitivity attenuates almost completely to zero in high spatial frequencies. The peak of its chromatic-CSF (in *fc*) is at the lowest spatial frequencies and its sensitivity drops to zero at the same pace as the human chromatic-CSF.

**Figure 4:**
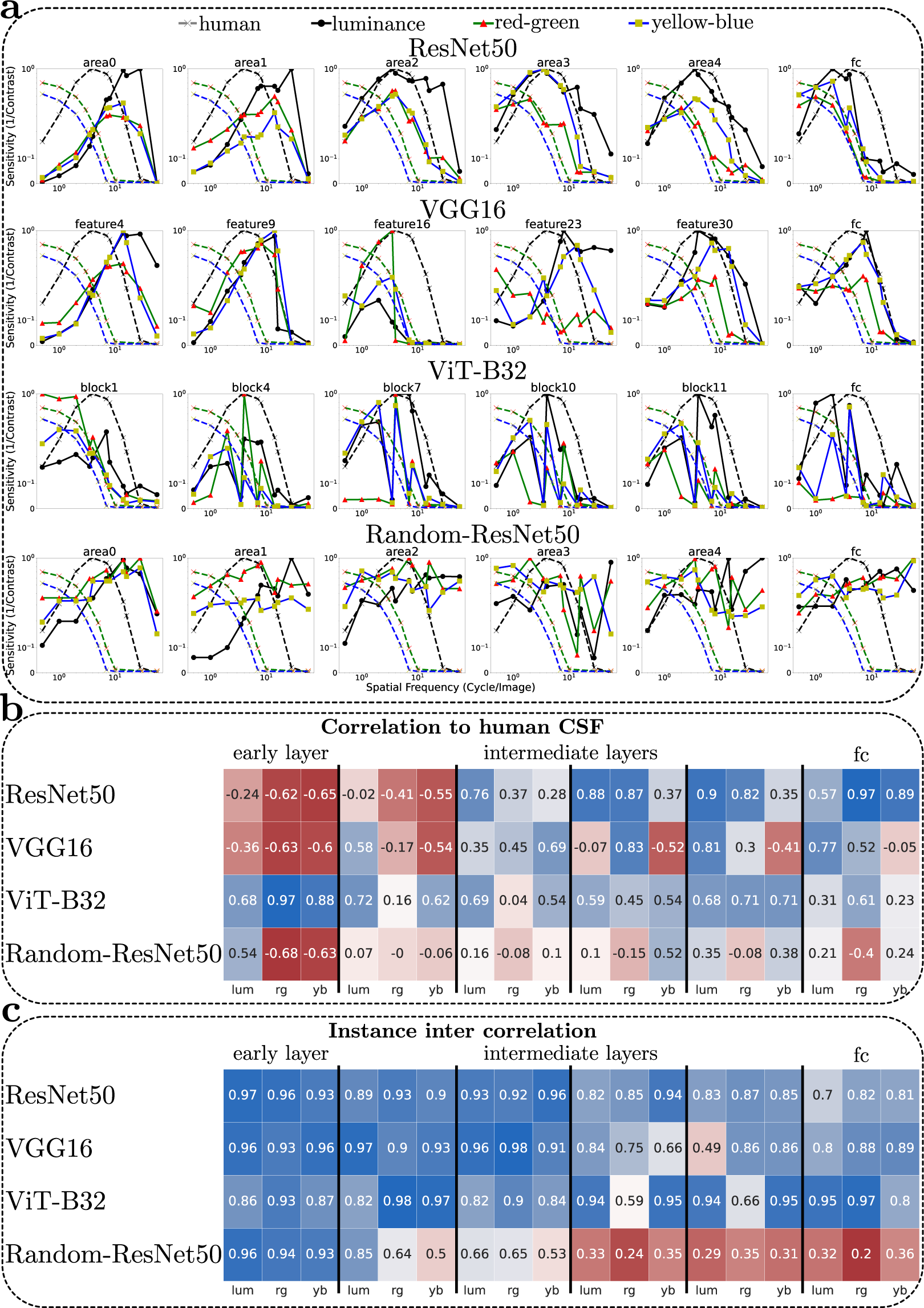
**a**: The CSF of three ImageNet pretrained networks and *Random-ResNet50* trained only on the contrast discrimination task. **b**: The correlation to the human CSF. Cells are colour-coded, blue indicates a higher correlation and red is lower. **c**: The average correlation coefficients across CSF of ten instances.

Interestingly, we also observe a striking difference across architectures in the depth of feature space (shallow, intermediate or deep layers) that explains the human CSF best:

1. While human-like luminance-CSF appears in intermediate layers of ResNet50 (i.e., *area2–4, r* = 0.90), the chromatic-CSF is more a late representation (e.g., the *fc* layer, *r_rg_* = 0.97 and *r_yb_* = 0.89), especially the yellow-blue channel. All channels jointly considered the *fc* layer best captured the human CSF.
2. While human-like luminance-CSF (*r* = 0.81) is a late representation in VGG16 (i.e., *feature30* and *fc*), the chromatic-CSF appears in intermediate layers, red-green in *feature23* (*r* = 0.83) and yellow-blue in *feature16* (*r* = 0.69).
3. In ViT-B32, human-like CSF is better captured in chromatic channels, particularly as an early representation (i.e., *block1, r_rg_*= 0.97 and *r_yb_* = 0.88). The luminance-CSF is relatively stable across all early to intermediate layers (*r* = 0.72) and it drops notably in the *fc* layer (*r* = 0.31).

Thus, the human-like CSF appears in shallow, intermediate, or deep layers across varying neural network architectures. Given the dimensionality of the feature space differs across layers and architectures (both in terms of the spatial resolution and the number of kernels/units), the spatial resolution of the internal representation cannot explain the results, in line with previous works on CSF of deep networks (Li et al., 2022). The activation maps for ResNet50 and VGG16 contain about the same number of pixels, and the feature space of Vit-B32 is always a two-dimensional vector of the same number of elements. This finding suggests the CSF is not necessarily a result of low-level feature tuning, and mid- and high-level features might well contribute to its shape.

### 3.3. Control experiments

To evaluate the importance of task-dependent features (from the pretrained network) versus the contrast-discriminator linear classifier in producing the CSF, we conducted a control experiment whereby we additionally trained a linear classifier on top of a ResNet50 with random weights (*Random-ResNet50*). If the linear classifier was the underlying cause of the networks’ CSF, we should also expect to obtain human-like CSF in this scenario. However, the CSF of the *Random-ResNet50* does not match the global shape or peak sensitivity of the human CSF (see Figure 4). Qualitatively, the luminance-CSF of Random-ResNet50 tends more towards a high-pass filter and its chromatic-CSFs do not show a large difference between the maximum and minimum sensitivities (i.e., resembling a flat sensitivity at all spatial frequencies). This dissimilarity is not due to the *Random-ResNet50*’s poor performance, as its classification accuracy in the training stage is on par with pretrained networks. These results suggest that the emerging CSF in pretrained networks are driven by a set of features that have been learnt during the pretrained task (i.e., object classification).

To examine the reliability of the pretrained features, we trained ten instances of the contrast-discriminator linear classifier on top of the same pretrained network. We computed the correlation between the CSF of ten instances using the leave-one-out cross-validation methodology. Figure 4-c reports the average correlation coefficients for four networks at six different layers. The obtained CSFs for ImageNet pretrained networks correlate highly among ten instances of linear classifiers suggesting a similar global shape of the CSF irrespective of the contrast-discriminator. The main difference is the amplitude of sensitivity that probably originates from the random contrast modulation of training images given all other factors are identical across instances, e.g., the frozen pretrained features and the weight initialisation of the linear classifier. Hence, it seems that instances with higher amplitude must have seen lower contrast images more frequently during their training leading to a more sensitive linear classifier.

In addition, we can observe that the correlation of CSFs is very low across ten instances of *Random-ResNet50* except in *area0*. The main difference among CSFs of *Random-ResNet50* is not the amplitude of sensitivity but rather the shape of the function, suggesting that unstable pretrained features allow the linear classifier to learn to do the task using different features.

We also investigated the impact of the linear classifier type. We trained six instances of a linear support vector machine (SVM) instead of a linear neural network (NN) on the output of ImageNet pretrained ResNet50. The SVM expects all inputs together instead of in small batches. Consequently, the memory of our hardware allowed us to input the SVM only with 1K samples (as opposed to 150K of NN). Nevertheless, the patterns of resulting CSFs across all areas remained very similar. The correlation coefficient between ten instances of NN and six instances of SVM is 0.76 (averaged across all layers and channels). The correlation is noticeably higher (*r* = 0.85) if we exclude the *fc* layer that requires more training samples due to its small feature size (a vector of 1000 elements, i.e., more than two orders of magnitudes smaller than all other areas).

Based on the above-conducted experiments, we can conclude the features of the pretrained networks establish the CSF of the pretrained network, not the linear classifier.

## 4. Visual tasks

In the previous section, we demonstrated that the human-like CSF appears in ImageNet pretrained networks that perform object recognition. In this section, we examine how the network’s task shapes its CSF using the Taskonomy dataset (Zamir et al., 2018), which contains about four million images (mainly indoor scenes from 2265 different buildings) and their corresponding labels for 25 computer vision tasks. The dataset also provides pretrained networks with an encoder-decoder architecture for all visual tasks^3^. The encoder is the same across all pretrained networks (i.e., ResNet50), which maps the input image (224 *×* 224) into a latent representation of size 1568 (14 *×* 14 *×* 8). The decoder design varies according to the dimensionality of the task output. The encoder offers a unique opportunity to study the role of visual tasks on the representation a network learns, given its architecture is identical across tasks and has been trained on the same set of images. Similar to ImageNet pretrained networks, we trained a linear classifier on top of the encoder’s extracted features from each Taskonomy pretrained network.

Figure 5-a shows the correlation coefficient between the human CSF and the Taskonomy pretrained networks across several layers. In *area0* many tasks tend to be negatively or weakly correlated to the human-like CSF. All tasks show very similar CSFs in *area0* (*r* = 0.95 across tasks, note the small standard error in Figure 5-b), whereas the CSF of other *areas* across tasks are weakly correlated (on average about 0.37, note the large standard errors in Figure 5-b). For several tasks, the networks’ CSF does not relate in the slightest to the human CSF (e.g., the pretrained networks on the *vanishing point* and *egomotion* tasks). This is not because these tasks are meaningless. In fact, both tasks are of great importance in importance in computer vision algorithms; *vanishing point* provides insights into the scene geometry by predicting the point on the perspective image plane where parallel lines converge, and *egomotion* defines the camera’s motion relative to the rigid scene. The dissimilarity of their CSF to the human CSF is rather because the features tuned to these tasks do not exhibit a strong preference for a specific spatial frequency or they are sensitive to spatial frequencies that humans are not.

**Figure 5:**
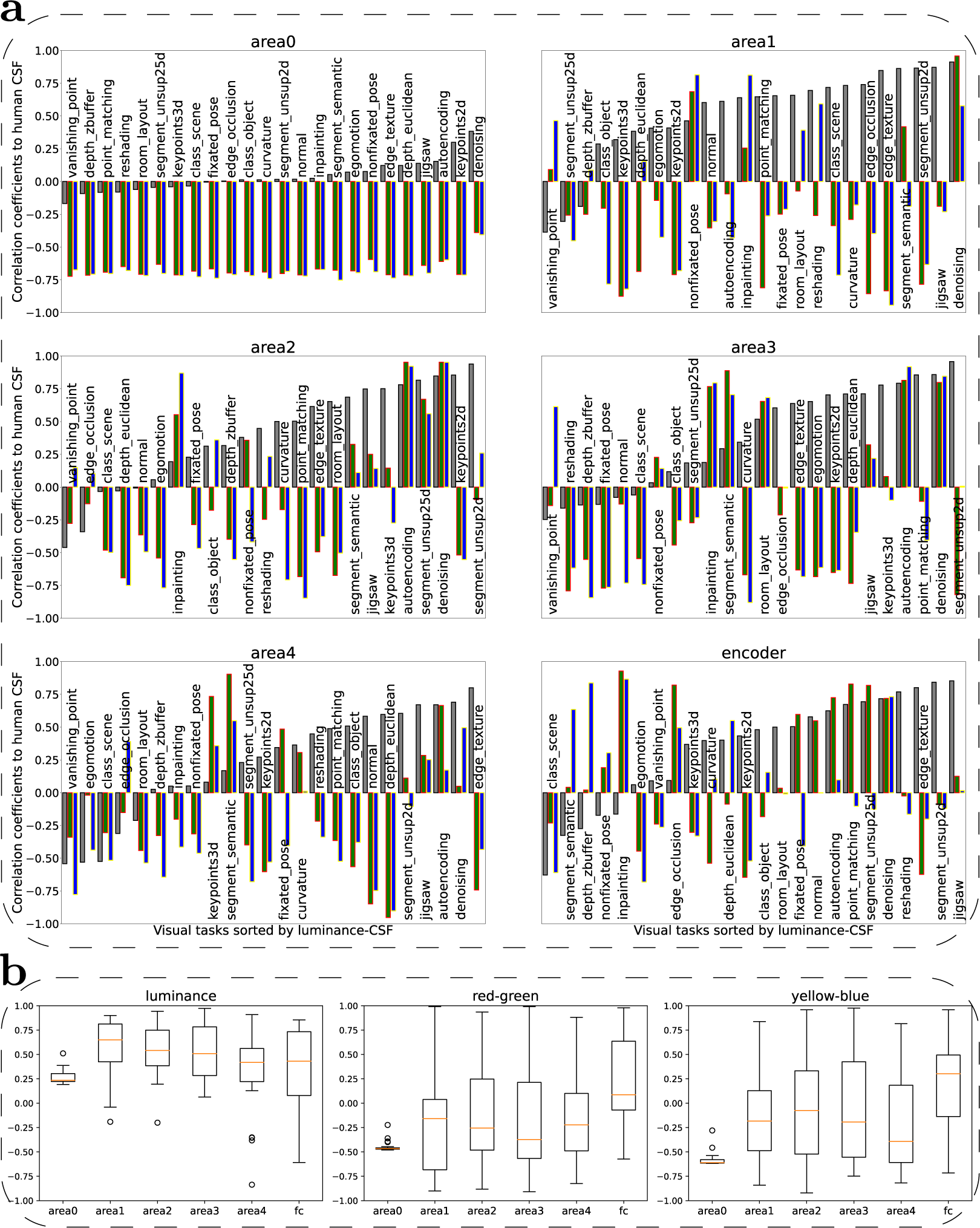
The results of Taskonomoy pretrained networks. **a**: The correlation of all Taskonomy pretrained networks to human CSF sorted by the correlation to the luminance CSF. Grey bars indicate the luminance-CSF, and green and blue bars are the chromatic-CSF (red-green and yellow-blue, respectively). **b**: The distribution of the correlation to human CSF for Taskonomoy networks. The box extends from the first to third quartile values of the data, with an orange line at the median. The vertical lines represent the most extreme, non-outlier data points.

Generally, the networks’ luminance-CSF correlates higher with human data (see Figure 5-b). Averaged across all tasks and areas, the correlation between networks and human CSF is 0.38 for the luminance channel and 0.15 for both chromatic channels. The single-task nature of these networks might explain this difference. An alternative explanation might be the convolution of the RGB channels in the networks’ first layer that collapses the three channels into a single stream of processing, which is unlike the human visual system that processes the chromatic channels in parallel up until the primary visual cortex. While there is a variation across tasks, generally, we observe that early to intermediate features better match the luminance-CSF (peaking at *area1*), and deeper features best explain the chromatic-CSFs (peaking at *encoder*).

### 4.1. Low-level tasks

Figure 6 illustrates the CSF of two Taskonomy pretrained networks trained on *denoising* and *autoencoding*. We consider these low-level because their goal has little direct relevance to real-world behaviours (i.e., they can be considered as a common preprocessing task for all visual tasks). The *autoencoding* network is optimised to find a low-dimensional latent representation of the input space, and the *denoising* network is optimised to obtain similar representations for similar inputs irrespective of the perturbation in the input. The *denoising* and *autoencoding* pretrained networks faithfully capture the human CSF in both luminance and chromatic channels (i.e., above 0.90 average correlation in *area2*). These findings match well previous reports explaining the relevance of low-level vision tasks in the emergence of human-like CSF (Li et al., 2022).

**Figure 6:**
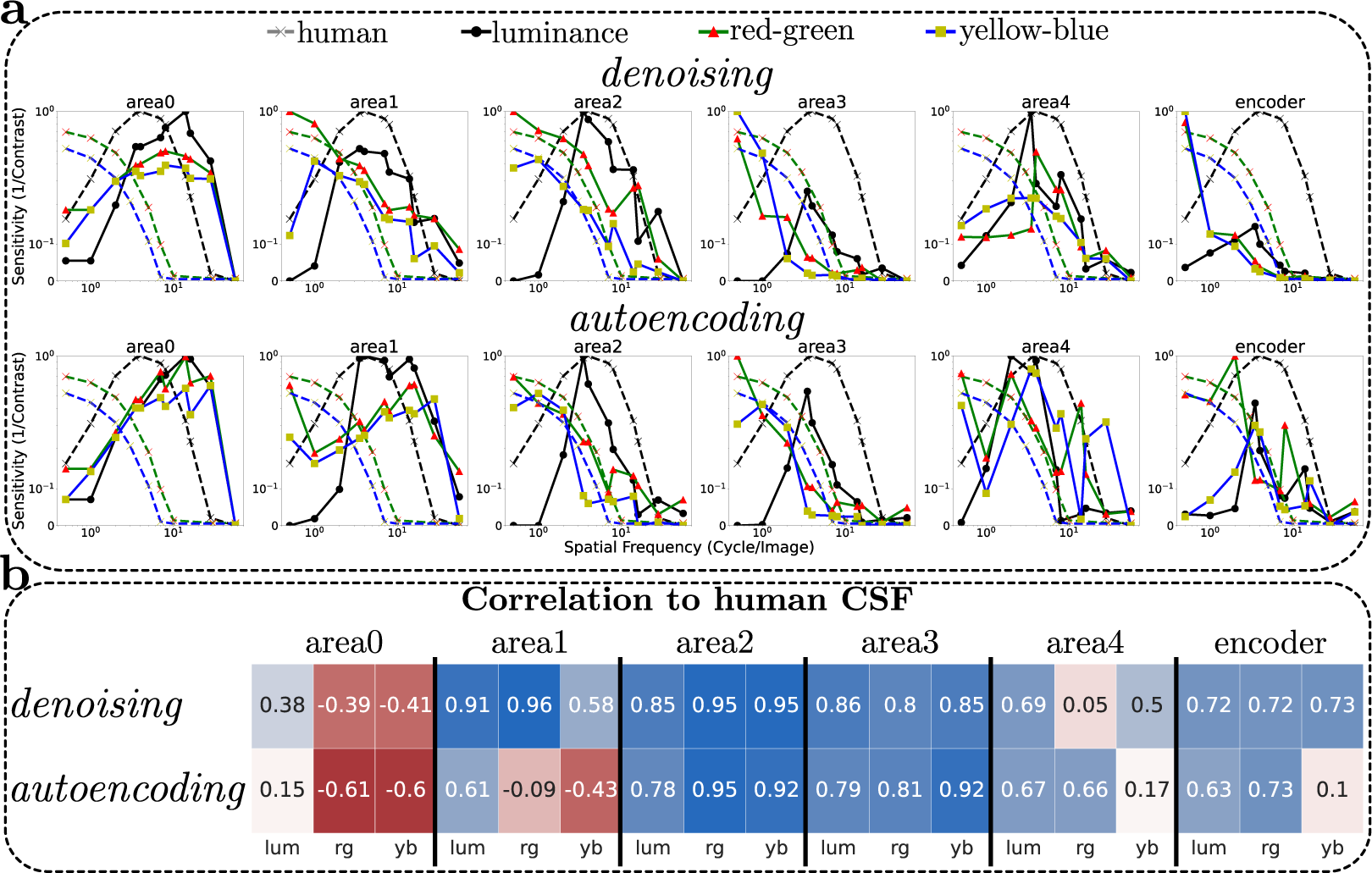
**a**: The CSF of two Taskonomoy pretrained networks. **b**: The correlation to the human CSF.

Features of other areas in *denoising* and *autoencoding* networks also match the shape of human CSF, for instance, the *area3* of both networks or *area1* in the denoising network. This is at odds with Li et al. (2022) that reported human-like CSF only emerges in shallow CNNs optimised on similar tasks, and deeper CNNs fail to obtain human-like CSF although they reach the quantitative goal better than shallower networks. Our results suggest deeper networks of *denoising* and *autoencoding* also capture the human CSF. Nevertheless, there is a small discrepancy in comparison to the human CSF. We often observe that the amplitude of chromatic-CSF is larger than the luminance-CSF in these two networks.

### 4.2. Mid-level tasks

While low-level tasks such as *denoising* and *autoencoding* best match the human CSF for both luminance and chromatic channels, several networks pretrained on mid-level visual tasks explain the luminance-CSF, see Figure 5-a; e.g., *edge texture* (extracting 2D edges of objects), *curvature* (measuring the surface bends in three directions), *keypoints2d* (detecting complex patterns that are invariant across multiple images such as corners and junctions), *normal* (estimating the surface normal in 3D), *point matching* (classifying if centres of two images match or not), etc. These mid-level tasks are also thought to underlie biological vision, e.g., edge detection (Marr, 1982), curvature processing (Yue et al., 2014), and corner detection (Tang et al., 2018). This co-occurrence suggests that mid-level features play an important role in shaping our CSF. Interestingly, the chromatic-CSF of all these mid-level tasks disagrees strongly with the human CSF. These results suggest that mid-level features significantly shape the luminance-CSF while playing little role in the chromatic-CSF.

In addition to the final goal of a task, the approach a system takes to solve that task impacts its corresponding features, and thus the resulting CSF. Formulating it in Marr’s levels of information-processing, the computational level influences the representation a system learns (Marr, 1982). This is noticeable in the depth estimation task for which two pretrained networks are available (*depth euclidean* and *depth zbuffer*). The Euclidean depth measures the distance from each pixel to the camera’s optical centre, whereas the Z-buffer depth measures the distance to the camera plane. Humans typically estimate depth using the Euclidean distance, whereas computer vision and computer graphics applications compute the Z-buffer depth (Zamir et al., 2018). Interestingly, human-like luminance-CSF only emerges in the pretrained network of *depth euclidean* and not the *depth zbuffer*, suggesting that deep networks trained on biologically plausible tasks are more representative of human visual behaviour (Neri, 2022).

### 4.3. High-level tasks

Human-like CSF also appears in Taskonomy pretrained networks performing high-level visual tasks (see Figure 5-a) such as scene segmentation and object classification, e.g., *segment semantic* (pixel-wise semantic labelling by distillation knowledge from COCO networks), *segment unsub25d* (unsupervised segmentation based on RGB-D-Normals-Curvature features), and *class object* (object recognition by distillation knowledge from ImageNet networks). The ImageNet experiment suggested that human-like CSF can emerge in different depths of visual processing depending on the network’s architecture (Figure 4). The Taskonomy experiments imply a similar phenomenon for different high-level tasks. For instance, the highest correlation to human CSF is obtained in *area4* of *segment semantic, area2* of *segment unsub25d*, and *encoder* of *class object* network.

We compared the segmentation versus classification task more thoroughly by training ten instances on the contrast-discriminator linear classifier on top of the Taskonomy pretrained networks. The results suggest that human-like CSF appears more regularly in the segmentation networks, and typically they better match the human CSF (i.e., averaged across all layers and channels *^r^_segmentation_* − *^r^_classification_* > 0.30).

Given, however, that the high-level Taskonomy tasks are trained to match ground truth labels derived automatically from state-of-the-art deep networks rather than human observers, the networks for the high-level Taskonomy tasks may show atypical features for segmentation and classification. Therefore, we decided to compare the semantic segmentation networks from the COCO dataset (Lin et al., 2014) to the ImageNet object recognition (i.e., humans have labelled the ground truth for both datasets).

We evaluated four pretrained semantic segmentation networks (DeepLab50, DeepLab101, FCN50, FCN101) that label each pixel of an image into twenty categories. DeepLab (Chen et al., 2018) and FCN (Shelhamer et al., 2017) encode the input images with a ResNet architecture (50 or 101, respectively), followed by a decoder that generates the segmentation maps. To facilitate a direct comparison to ImageNet networks, we trained the contrast-discriminator linear classifier on top of the encoder features. Figure 7 reports the CSF of COCO pretrained networks averaged over four instances. These results match those of Taskonomy networks suggesting segmentation tasks better capture the human CSF in comparison to classification tasks (i.e., averaged across all layers and channels *r_segmentation_ − r_classification_ >* 0.10). The underlying representational difference between these two seemingly similar tasks might be originated from the spatial resolution of networks’ output space, in which the segmentation is pixel-wise while classification is a single label for the entire image.

**Figure 7:**
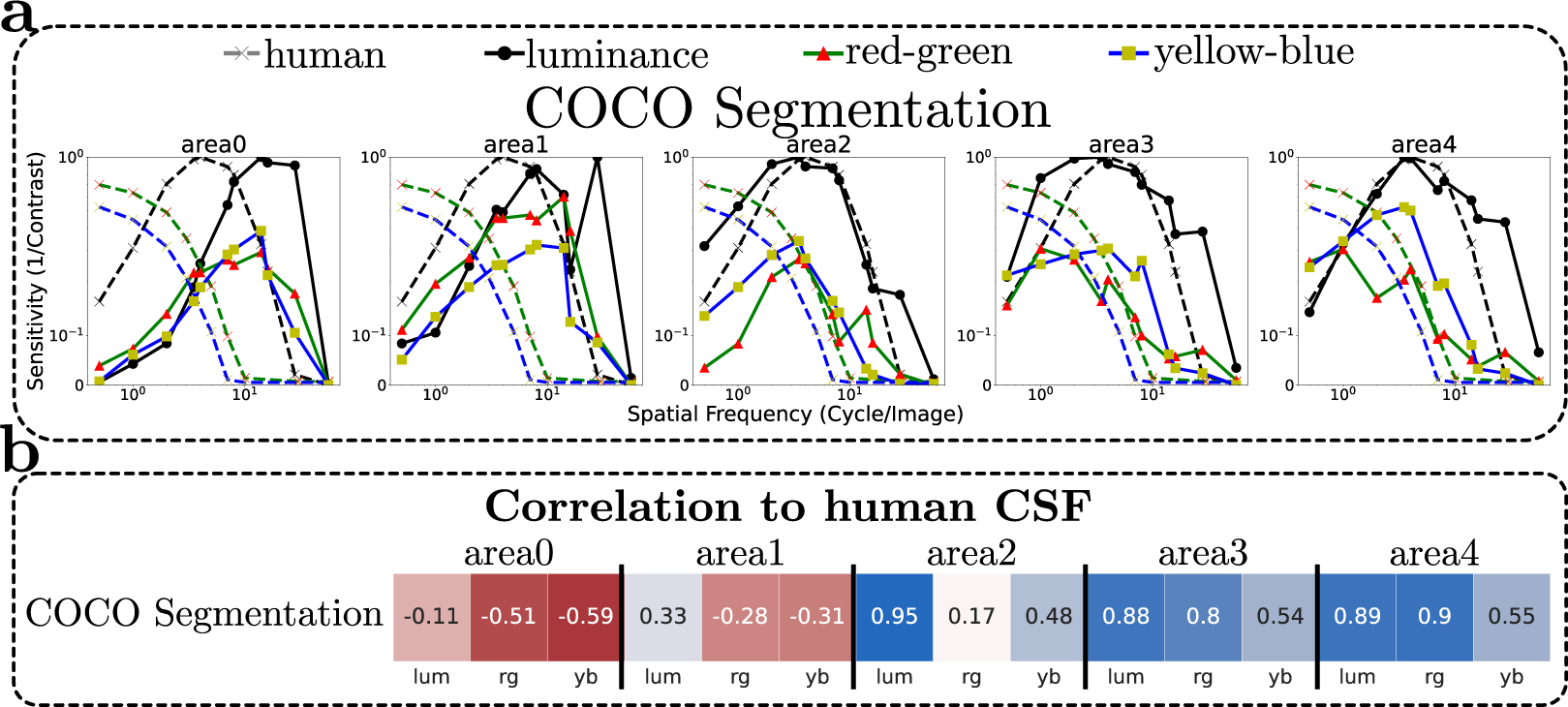
**a**: The CSF of semantic segmentation networks pretrained on the COCO dataset (averaged over DeepLab50, DeepLab101, FCN50, and FCN101). **b**: The correlation to the human CSF.

### 4.4. Structure of tasks

We calculated the correlations of the CSFs among all the pairs of visual tasks and used that as a similarity matrix to analyse the structure of the tasks. Figure 8 shows the correlation between a pair of network CSF averaged over five intermediate layers (*area0* is excluded given its high degree of correlation across all visual tasks). We clustered them using the UPGMA, unweighted pair group method with arithmetic mean, hierarchical clustering method (Müllner, 2013). In the luminance channel, we can observe three clusters. (1) The human CSF is clustered with a number of pretrained Taskonomy networks from different levels of visual tasks, low (e.g., *denoising* and *autoencoding*), mid (e.g., *jigsaw* and *edge texture*) and high (e.g., *segment unsup2d*). (2) A small group of tasks (e.g., *nonfixated pose, egomotion* and *class scene*) have the peak of their CSF in high spatial frequencies. (3) Many tasks (e.g., *room layout, curvature, depth euclidean*, etc.) have the peak of their CSFs in middle spatial frequencies, therefore partially correlating to both other groups.

**Figure 8:**
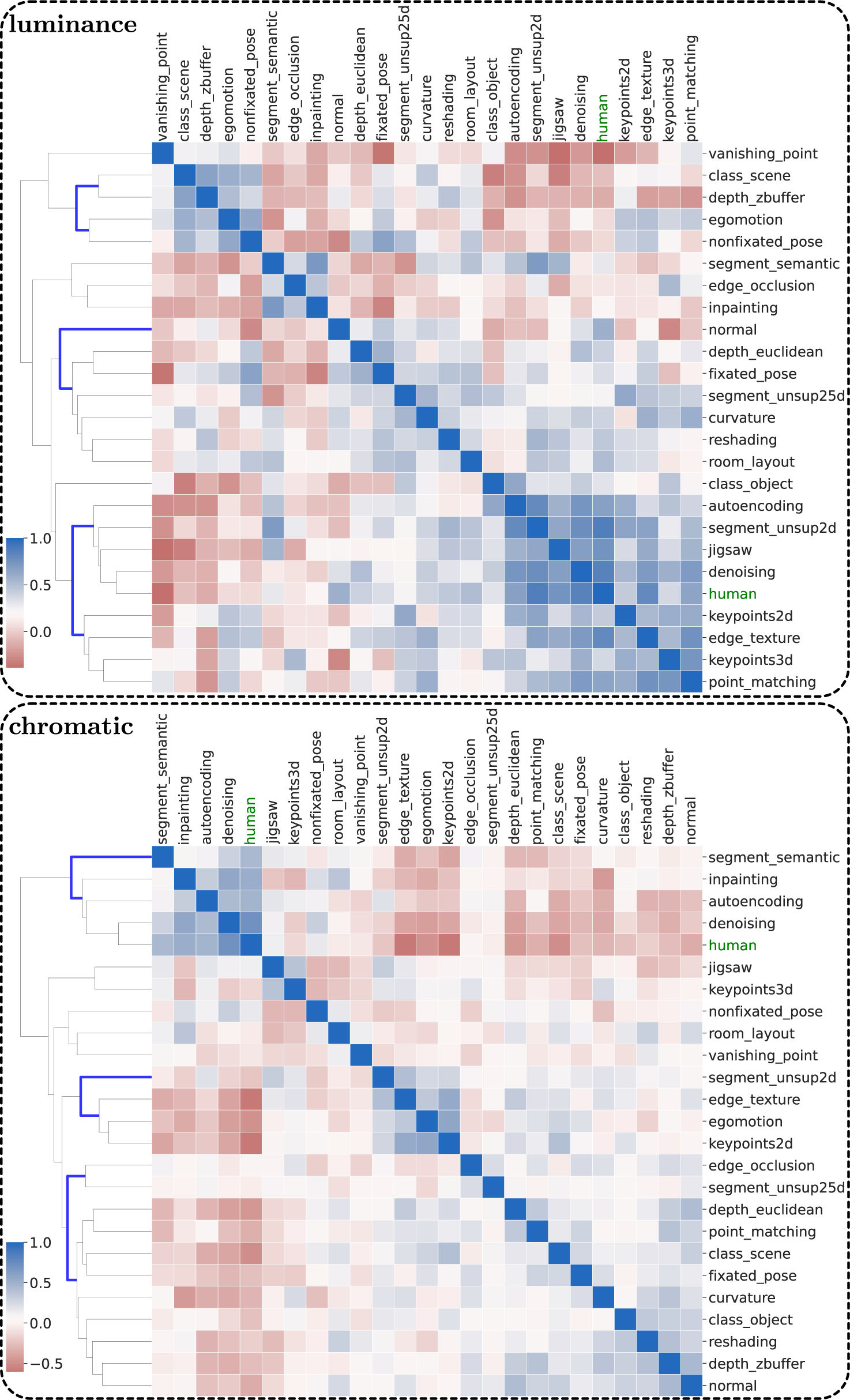
Hierarchical clustering of networks according to their degree of similarity in luminance(top) and chromatic-CSFs (bottom). The coefficients are averaged over all areas excluding *area0* where all networks are highly correlated.

This analysis reiterates that a system’s approach to solving a task impacts its corresponding features. For instance, *class scene* and *class object* are conceptually very similar tasks. They both predict the category of the input image among a predefined set of classes. Interestingly, the CSF of *class scene* peaks at higher spatial frequencies, while the peak of the sensitivity for *class object* is at a lower spatial frequency, therefore better matching the human CSF. Intuitively, to solve the task of scene classification a given visual system has to rely on a collection of a finer set of features to distinguish between, for example, a picture of a living room and an office. Contrary to that, to solve the task of object classification a given visual system has to rely on a coarser set of features (e.g., the global shape of an object) to distinguish between, for example, a picture of a television and a sofa.

In the chromatic channels, we can also discuss the obtained clusters within three major groups. (1) The human CSF is grouped with four tasks, namely, *denoising, autoencoding, inpainting* and *segment semantic*. (2) A large set of tasks are clustered together mainly because their chromatic-CSF is relatively flat (e.g., *normal, depth zbuffer, curvature, point matching*, etc.), essentially their chromatic tuning is not strongly influenced by the spatial frequency. (3) The third group (*segment unsup2d, edge texture, egomotion* and *keypoints2d*) shows more a broad-band chromatic-CSF with a peak at middle spatial frequencies.

## 5. Vision-language networks

Recently vision-language models like CLIP (Contrastive Language-Image Pretraining) have achieved tremendous success in zero-shot learning and as the backbone for image-generating networks. Therefore, we extended our investigation of CSF in deep networks to two versions of the vision-language models, *CLIP ViT-B32* and *CLIP ResNet50* (Radford et al., 2021). Both models have a transformer text encoder and an image encoder (i.e., modified versions of ViT-B32 and ResNet50, respectively) that are jointly optimised to predict the correct pairings of a batch of image text. We trained a contrast-discriminator linear classifier on top of features extracted from their image encoder that are the same architectures for ImageNet pretrained networks. Figure 9 illustrates the corresponding CSFs averaged over ten instances (the average instance inter-correlation is 0.85 for *CLIP ViT-B32* and 0.75 for *CLIP ResNet50*. The results suggest that human-like CSF appears in several layers of *CLIP ViT-B32*. While the luminance-CSF is best captured in intermediate to deep layers (*r* = 0.92), human-like chromatic-CSF emerges more in early to intermediate layers (*r* = 0.94). Contrary to this, in *CLIP ResNet50* human-like CSF only appears in the luminance channel of two intermediate layers (with a lower correlation *r* = 0.71). It was previously reported that *CLIP ViT-B32* scores closely to humans across several psychophysical experiments, specifically in error consistency (Geirhos et al., 2021), which is in agreement with our findings of CSF in this network. While they do not report the results of *CLIP ResNet50*, it can be argued that error consistency of *CLIP ViT-B32* to human data cannot be explained merely by the task/loss/dataset, otherwise, *CLIP ResNet50* should have obtained more human-like CSFs.

**Figure 9:**
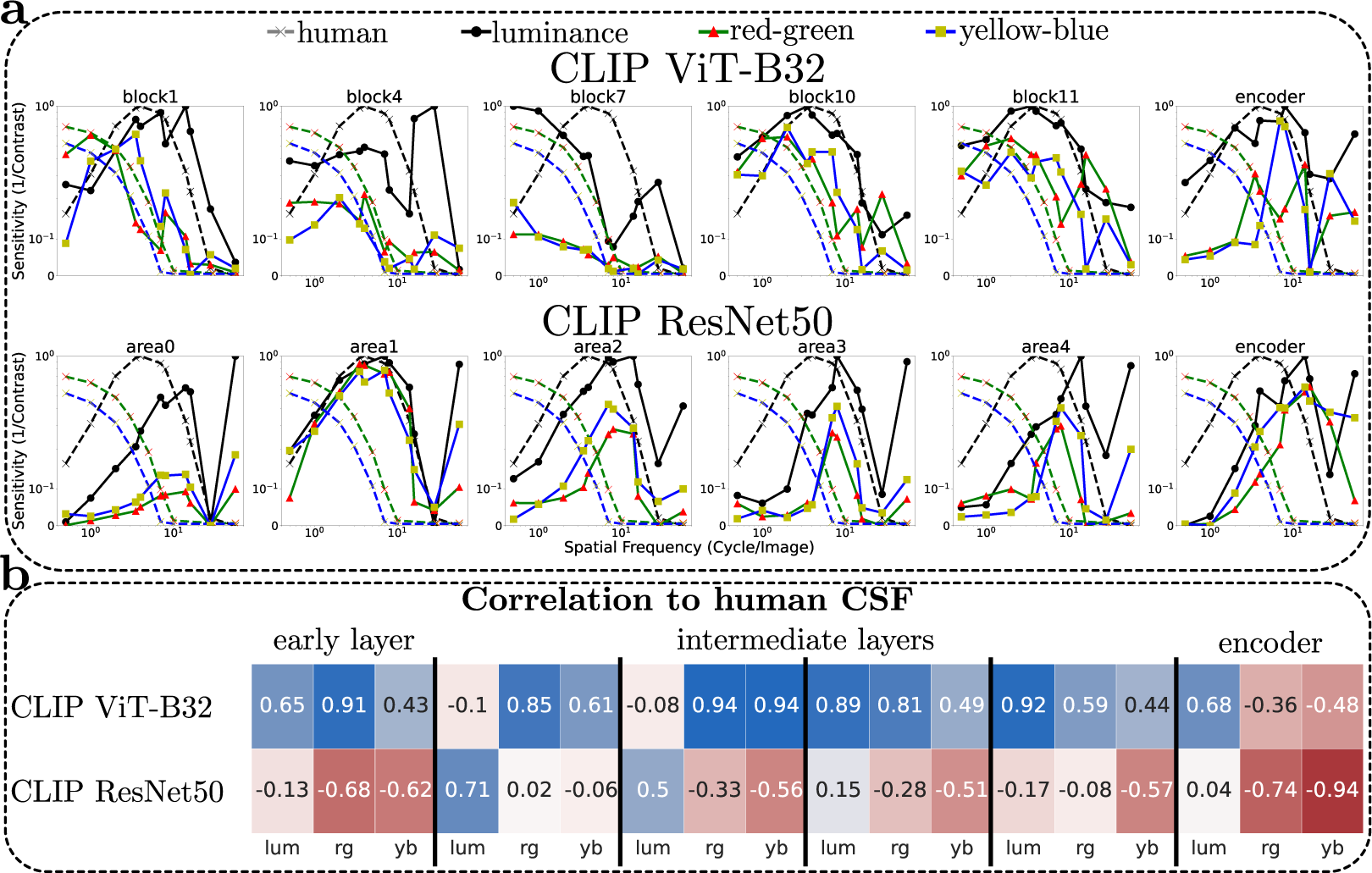
**a**: The CSF of CLIP vision-language models with two different image encoders. **b**: The correlation to the human CSF.

## 6. Discussion

The contrast sensitivity function (CSF) measures the visibility threshold of sinusoidal gratings as a function of spatial frequency. We measured the CSF of deep neural networks using the same paradigm as human psychophysics. The CSF of many pretrained networks exhibited the characteristics of the human CSF, a band-pass inverted-U shape function in the luminance channel, and two low-pass filters of similar properties in chromatic channels. This finding suggests the relative significance of spatial frequencies in static images is alike in the human visual system and DNNs that perform meaningful visual tasks. Contrary to previous works that have attributed the shape of human CSF mainly to low-level explanations (Graham, 1972; Wandell, 1995; Atick, 1992; Li et al., 2022; Gomez-Villa et al., 2020) like the centre-surround mechanism of ganglion cells, our results hint toward the contribution of features from all levels of visual hierarchy to the tuning shape of DNNs’ CSF.

ImageNet experiments show that human-like CSF can appear at distinct depths of visual processing depending on the network’s architecture (i.e., early, intermediate and deep layers in different architectures capture the human luminance- and chromatic-CSFs). While the exact internal representation of these object recognition networks is not fully known, they share common properties in their hierarchy of visual processing, such as kernels in earlier layers responding to oriented bars, intermediate layers to textural patterns, and deeper layers to object parts (Lindsay, 2020). Therefore, the emergence of human-like CSF at several depths of different architectures suggests the underlying representation driving the shape of human CSF is not merely limited to low-level visual features. The Taskonomy experiments strengthen this observation by demonstrating that human-like CSF can appear in pretrained networks of an identical architecture optimised for several tasks with different levels of visual abstraction. Furthermore, the exact area best capturing the human CSF in Taskonomoy pretrained networks does not indicate a bias towards low-level features. For instance, the human CSF is captured best by *area2* of *denoising* (i.e., an intermediate layer of a lowlevel task network) and the *area4* of *segment semantic* (i.e., a deep layer of a high-level task network). If low-level visual features modulated the CSF, we would expect an inverse pattern of results for the segmentation network, i.e., human-like CSF would appear more in earlier layers of a segmentation network that presumably represent low-level visual features.

While the examined networks matched well the shape of human CSF, they differ in their absolute sensitivity in two aspects, (i) the magnitude of the contrast sensitivity is substantially larger in deep networks, and (ii) the relative difference of magnitude between the luminance and chromatic channels is moderately larger in humans. Several factors influence the absolute sensitivity of a system. The sensitivity of the sensory system (photoreceptors) is the starting point. A system input with a single pixel in the RGB space maximally obtains a 255 sensitivity magnitude (*uint8* precision). The examined networks are input with RGB images, but their architecture space is floating precision. Thus, although the kernels operating on the input pixels are exposed maximally to 0.392% (^1^) contrast, the upstream kernels have access to a higher contrast precision in the internal feature space. A theoretical kernel with a discrimination power on the entire spatial resolution of input across all three channels has access to a 38*M* (224 *×* 224 *×* 3 *×* 255) granularity contrast. This line of thinking can potentially explain the hypersensitivity of the studied networks. Nevertheless, the maximum sensitivity we observed in the Taskonomy pretrained networks was a more moderate figure (52*K*) occurring in the first layer of ResNet50 (*area0*), and the maximum sensitivity in other areas is about 20*K* with an average value (500) that is not far away from the human sensitivity.

The absolute sensitivity of a system essentially refers to its accuracy (i.e., it is defined as the inverse of the contrast in the psychometric function where the accuracy is 75%). Therefore, networks with higher sensitivities are more accurate at test time (i.e., input with sinusoidal gratings). We investigated whether these networks also obtain higher accuracy in natural images (i.e., the contrast discrimination training). Nevertheless, we did not find any relationship between the training time accuracy and the network’s sensitivity, suggesting that the hypersensitivity of some networks originates from their internal representation irrespective of their CSF shape.

The human contrast sensitivity is lower in both chromatic channels in comparison to the luminance one when expressed in RGB space. We see a similar phenomenon in deep networks whose luminance sensitivity is higher but with a smaller absolute difference in the chromatic channels. The low sensitivity of the yellow-blue channel in the human visual system is explained by the lack of short photoreceptors in the fovea (Stromeyer et al., 1978; Williams and Collier, 1983). Nevertheless, the same reasoning falls short in explaining the red-green sensitivity threshold. Therefore, photoreceptors alone cannot explain the difference in amplitude of chromatic- and luminance-CSFs. Furthermore, the mosaic distribution of RGB photoreceptors in digital cameras is vastly different from the distribution of cone photoreceptors in the human retina (Ramanath and Snyder, 2003), therefore, the similarity in their chromatic-CSFs, i.e., no attenuation in low spatial frequencies (Mullen, 1985), must originate from some other visual features common to both systems.

Among several interesting questions that one can wonder about is the phenomenon of noise. The efficiency of signal discrimination in the human visual system is often defined as a function of stimuli noise (Burgess et al., 1981). Consequently, many studies comparing the discrimination threshold of DNNs to humans introduce noise to the network’s input signal (Srivastava et al., 2022) arguing that in noise-free settings even a naive system can perfectly discriminate between stimuli’ presence (e.g., our positive contrast sinusoidal gratings) and absence (e.g., our zero contrast uniform background). Indeed, in our experiments, the activation map of networks is always zero when the stimuli contrast is zero. Effectively, a rudimentary system comparing the average of absolute intensities can perfectly discriminate between the two stimuli up to the precision of floating points irrespective of the spatial frequency. In other words, the CSF of such a system would be a perfectly flat horizontal line. Nevertheless, the linear classifiers trained with natural images to identify the image with higher contrast cannot pick on this clue, as a result, the networks’ CSFs are never flat, demonstrating the inherent bias of pretrained networks for different spatial frequencies. It is worth noting that the human CSF is also measured with noise-free stimuli.

### 6.1. Extended CSF

Here, we extensively analysed the role of the network’s architecture and task on its contrast sensitivity function in static images. Yet another fundamental factor is the environment in which a visual system inhabits. The networks we examined were trained on datasets of natural images captured with commercial cameras (i.e., ImageNet, COCO, Taskonomy, CLIP). A systematic change of the training dataset for the pretrained task can answer how the environment shapes the system’s CSF. For instance, would training the same architecture and task (e.g., ResNet50 on object recognition) on aerial images results in a CSF whose peak lies on high spatial frequencies in line with the CSF of many large birds of prey like Falcon and Eagle (Hirsch, 1982)? Our preliminary results on filtered images of ImageNet support this hypothesis (Akbarinia et al., 2021).

The CSF of biological systems also depends on other factors, like the temporal frequency (Kelly, 1979), luminance level (Wuerger et al., 2020), and the interaction between them (Díez-Ajenjo and Capilla, 2010). Interesting questions remain on how such spatiotemporal-chromatic settings underlie more human-like CSFs in deep networks, where previous work hints that spatiotemporal aspects emerge in convolutional autoencoders performing low-level visual tasks of retinal noise and optical blur removal (Li et al., 2022).

Last but not least, one of the biggest challenges in neuroscience is understanding how single-neuron responses pool with other neurons yielding systems-level behavioural responses. Hence, it would be interesting to investigate how the response of individual kernels to stimulus contrast mediates the network’s CSF. Two ways of achieving this may be by (1) lesioning one kernel at a time and measuring the network’s CSF, and (2) computing the CSF of individual kernels by recording their activation to stimuli that vary in their contrast. These experiments would explain how the features at each kernel/layer/block influence the shape of CSF and its underlying mechanism.

## 7. Conclusion

Our work demonstrates that one can systematically conduct psychophysical experiments with artificial neural networks. We used a linear classifier to probe the internal representation of several pretrained networks spanning tasks and architectures. We validated the reliability of this approach in three control experiments and by showing a high degree of similarity across many instances. The instrumental linear classifier we used is not limited to measuring the contrast sensitivity function of a network, and it can be used for other psychophysics aiming to achieve a more direct comparison between artificial and biological systems (Bowers et al., 2022).

The pretrained networks exhibiting a human-like CSF are physically very distinct from the biological visual systems. In Marr’s terminology, the implementation level in artificial kernels and biological neurons differs substantially. Therefore, the similarity in the CSF of these two systems originates from the problem they try to solve (e.g., making sense of visual information). We observed in the Taskonomy experiments that task demand is crucial in the emergence of human-like CSF. While all Taskonomy pretrained networks are optimised to a meaningful visual task (with the same set of images), only a subset of those (mainly ecologically plausible tasks) turn out to closely match the human CSF. Similarly, biological vision selects and processes certain visual stimuli while ignoring others by allocating its limited resources to appropriately meet the demands of a particular task (Barlow et al., 1961). Thus, given that the spatial frequency information most important for several of these single-task networks coincides with those most sensitive to human observers, we suggest that the exact shape of our CSF is tuned to accommodate the needs of multiple tasks it must perform in the physical world.

## Acknowledgement

This study was funded by Deutsche Forschungsgemeinschaft SFB/TRR 135 (grant number 222641018) TP C2. This article has been presented in the form of an abstract at the Vision Science Society 2021 (Akbarinia et al., 2021).

## Appendix A Chromatic-gratings

Figure A.1 visualises this issue with an example of a 50% red-green contrast, which yields about 100% contrast in the R-channel, about 10% in the G-channel and 1% in the B-channel. Contrary to this, contrast modulation in the YP*_b_*P*_r_* colour space results in sinusoidal gratings of identical waves with different phases for the R- and G-channels (Mullen, 1985). Additionally, in the case of high-contrast gratings, the transformation from DKL to RGB might go beyond the RGB gamut. While this is practically not an issue for our purpose (we are dealing with very low-contrast stimuli), this problem does not exist for YP*_b_*P*_r_* as the coefficients of RGB channels are *±*1. Altogether, we report the YP*_b_*P*_r_* results in this manuscript to avoid any undesired artefacts. Nonetheless, we have also evaluated all our networks in the DKL, reaching an almost identical pattern of results (materials available on our GitHub https://github.com/ArashAkbarinia/DeepCSF).

## Appendix B ImageNet pretrained networks

Figure B.1 reports the CSF of three architectures at six different layers. We chose these three architectures because they are the extrema of the ImageNet pretrained networks (i.e., RegNet obtained the most humanlike luminance-CSF among ImageNet pretrained networks, ConvNeXt the worst and ViT-L32 the best chromatic-CSF), refer to Section 3.2 of the main text for more information about the networks’ internal representation. Here, also we observe considerable differences among architectures. While RegNet luminance-CSF greatly resembles the human CSF in the deepest layer (i.e., DKL YP*_b_*P*_r_ fc, r* = 0.94, the object classification feature space), its chromatic-CSF is less similar to the human’s, and more importantly, it only appears in the shallowest layer (*stem, r_rg_*= 0.79 and *r_rg_* = 0.59). Although ConvNeXt-Tiny captures the human CSF for the luminance channel in the layer *feature7* (*r* = 0.67) and the yellow-blue channel in *feature2* (*r* = 0.83), its CSF in the reg-green chromatic channel correlates only slightly to the human CSF (*r* = 0.30). Nevertheless, there is a high chance that other internal layers, whose CSF we did not compute, obtain higher correlations. ViT-L32 (a vision transformer architecture) obtains human-like chromatic-CSF in its earliest and deepest layers (*block1* and *fc, r ≥* 0.93) and luminance-CSF to a lower degree in its intermediate layers (*block13, r* = 0.58).

---

**Figure A.1:**
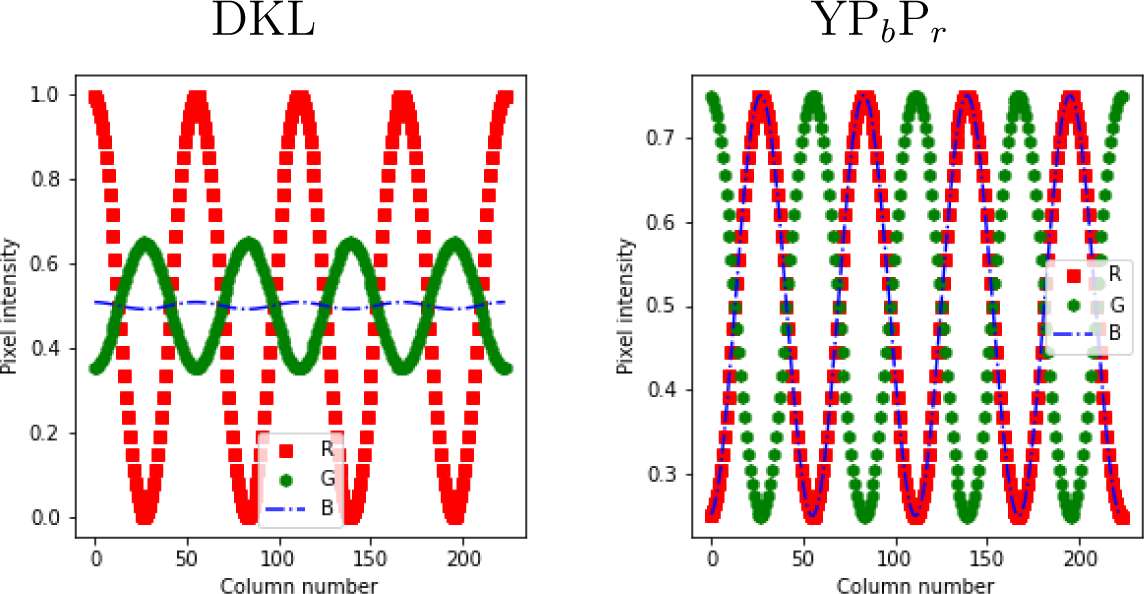
The RGB Profile of a sinusoidal grating of 50% red-green contrast.

**Figure B.1:**
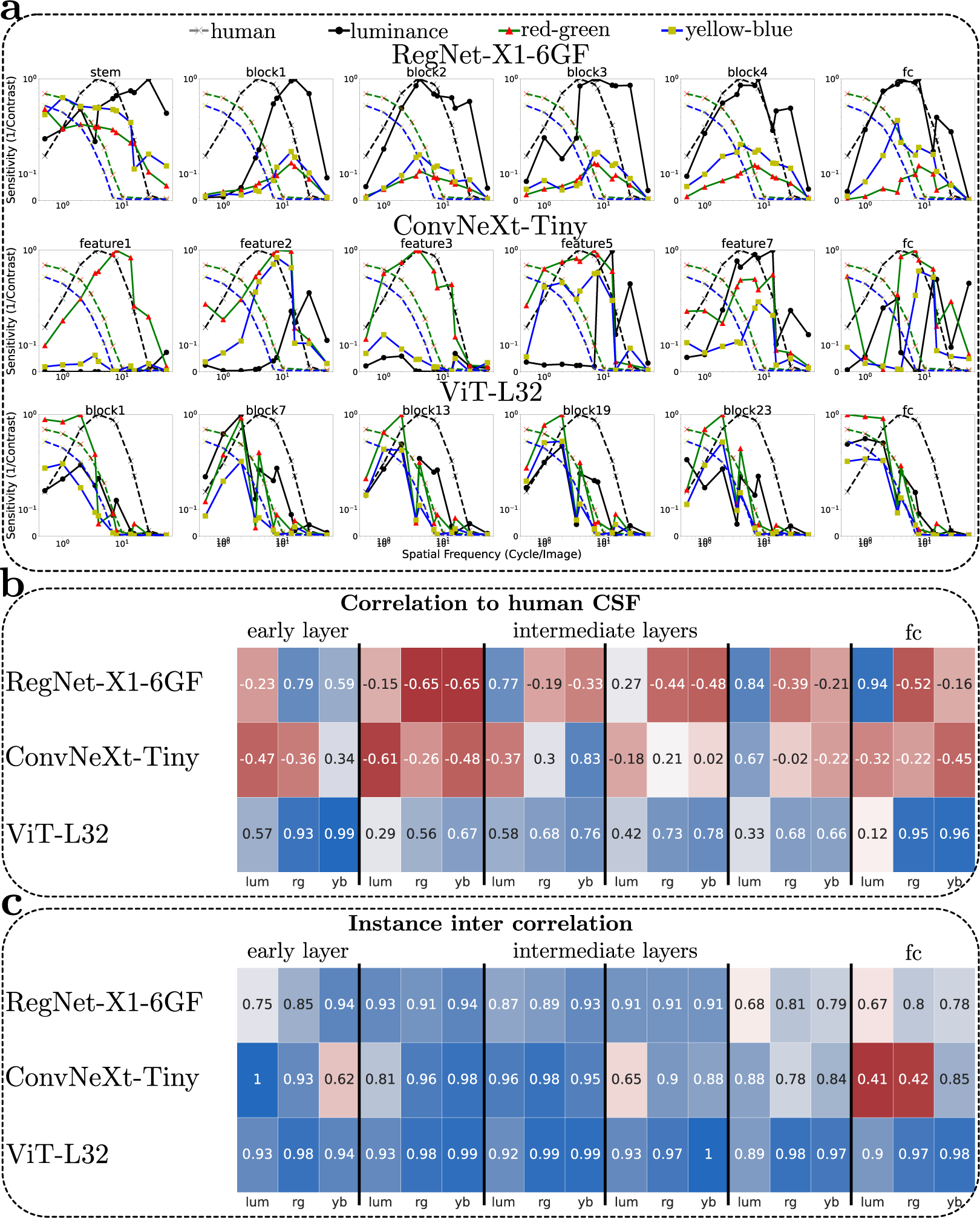
**a**: The CSF of three ImageNet pretrained networks averaged across ten instances. **b**: The correlation to the human CSF. Cells are colour-coded, blue indicates a higher correlation and red is lower. **c**: The average correlation coefficients across CSF of ten instances.

## Footnotes

1 All the experimental materials including the source code and weights of the examined networks are available on our project page https://arashakbarinia.github.io/projects/deepcsf/

2 https://pytorch.org/vision/stable/models.html

3 https://github.com/alexsax/midlevel-reps/

## References

Akbarinia, A., Gil-Rodriguez, R., 2020. Deciphering image contrast in object classification deep networks. Vision Research 173, 61–76.

Akbarinia, A., Gil-Rodŕıguez, R., 2021. Color conversion in deep autoencoders. Journal of Perceptual Imaging, 20401–1–20401–10.

Akbarinia, A., Morgenstern, Y., Gegenfurtner, K.R., 2021. Contrast sensitivity is formed by visual experience and task demands. Journal of Vision 21, 1996–1996.

Alain, G., Bengio, Y., 2017. Understanding intermediate layers using linear classifier probes, in: International Conference on Learning Representations.

Anscombe, F.J., 1973. Graphs in statistical analysis. The American Statistician 27, 17–21.

Atick, J.J., 1992. Could information theory provide an ecological theory of sensory processing? Network: Computation in neural systems 3, 213–251.

Atick, J.J., Li, Z., Redlich, A.N., 1992. Understanding retinal color coding from first principles. Neural computation 4, 559–572.

Atick, J.J., Redlich, A.N., 1992. What does the retina know about natural scenes? Neural computation 4, 196–210.

Barlow, H.B., et al., 1961. Possible principles underlying the transformation of sensory messages. Sensory communication 1, 217–234.

Barten, P.G., 1999. Contrast sensitivity of the human eye and its effects on image quality. SPIE press.

Bashivan, P., Kar, K., DiCarlo, J.J., 2019. Neural population control via deep image synthesis. Science 364, eaav9436.

Bisti, S., Maffei, L., 1974. Behavioural contrast sensitivity of the cat in various visual meridians. The Journal of Physiology 241, 201–210.

Bowers, J.S., Malhotra, G., Dujmovíc, M., Montero, M.L., Tsvetkov, C., Biscione, V., Puebla, G., Adolfi, F., Hummel, J.E., Heaton, R.F., et al., 2022. Deep problems with neural network models of human vision. Behavioral and Brain Sciences, 1–74.

Burgess, A., Wagner, R., Jennings, R., Barlow, H.B., 1981. Efficiency of human visual signal discrimination. Science 214, 93–94.

Cadieu, C.F., Hong, H., Yamins, D.L., Pinto, N., Ardila, D., Solomon, E.A., Majaj, N.J., DiCarlo, J.J., 2014. Deep neural networks rival the representation of primate it cortex for core visual object recognition. PLoS computational biology 10, e1003963.

Campbell, F., Green, D., 1965. Optical and retinal factors affecting visual resolution. The Journal of physiology 181, 576–593.

Campbell, F., Robson, J.G., 1968. Application of fourier analysis to the visibility of gratings. The Journal of physiology 197, 551–566.

Carandini, M., Heeger, D.J., 2012. Normalization as a canonical neural computation. Nature Reviews Neuroscience 13, 51–62.

Carney, T., Klein, S.A., Tyler, C.W., Silverstein, A.D., Beutter, B., Levi, D., Watson, A.B., Reeves, A.J., Norcia, A.M., Chen, C.C., Makous, W., Eckstein, M.P., 1999. Development of an image/threshold database for designing and testing human vision models, in: Human Vision and Electronic Imaging IV, SPIE. pp. 542–551.

Chen, L.C., Zhu, Y., Papandreou, G., Schroff, F., Adam, H., 2018. Encoderdecoder with atrous separable convolution for semantic image segmentation, in: Proceedings of the European Conference on Computer Vision, pp. 801–818.

Cichy, R.M., Khosla, A., Pantazis, D., Torralba, A., Oliva, A., 2016. Comparison of deep neural networks to spatio-temporal cortical dynamics of human visual object recognition reveals hierarchical correspondence. Scientific reports 6, 1–13.

Cornsweet, T., 1970. Visual perception. Academic press.

De Valois, R.L., Morgan, H., Snodderly, D.M., 1974. Psychophysical studies of monkey vision-iii. spatial luminance contrast sensitivity tests of macaque and human observers. Vision research 14, 75–81.

Deng, J., Dong, W., Socher, R., Li, L.J., Li, K., Fei-Fei, L., 2009. Imagenet: A large-scale hierarchical image database, in: IEEE Conference on Computer Vision and Pattern Recognition, pp. 248–255.

Díez-Ajenjo, M.A., Capilla, P., 2010. Spatio-temporal contrast sensitivity in the cardinal directions of the colour space. a review. Journal of Optometry 3, 2–19.

Eickenberg, M., Gramfort, A., Varoquaux, G., Thirion, B., 2017. Seeing it all: Convolutional network layers map the function of the human visual system. NeuroImage 152, 184–194.

Flachot, A., Akbarinia, A., Schütt, H.H., Fleming, R.W., Wichmann, F.A., Gegenfurtner, K.R., 2022. Deep neural models for color classification and color constancy. Journal of Vision 22, 17–17.

Geirhos, R., Narayanappa, K., Mitzkus, B., Thieringer, T., Bethge, M., Wichmann, F.A., Brendel, W., 2021. Partial success in closing the gap between human and machine vision, in: Proceedings of the Conference on Neural Information Processing Systems, pp. 23885–23899.

Geirhos, R., Temme, C.R., Rauber, J., Schütt, H.H., Bethge, M., Wichmann, F.A., 2018. Generalisation in humans and deep neural networks, in: Proceedings of the Conference on Neural Information Processing Systems, p. 7549–7561.

Geisler, W.S., 2008. Visual perception and the statistical properties of natural scenes. Annu. Rev. Psychol. 59, 167–192.

Gomez-Villa, A., Martín, A., Vazquez-Corral, J., Bertalmío, M., Malo, J., 2020. Color illusions also deceive cnns for low-level vision tasks: Analysis and implications. Vision Research 176, 156–174.

Graham, N., 1972. Spatial frequency channels in the human visual system: Effects of luminance and pattern drift rate. Vision research 12, 53–68.

Graham, N.V.S., 1989. Visual pattern analyzers. Oxford University Press.

Harmening, W.M., Nikolay, P., Orlowski, J., Wagner, H., 2009. Spatial contrast sensitivity and grating acuity of barn owls. Journal of Vision 9, 13–13.

Hashemi, H., Khabazkhoob, M., Jafarzadehpur, E., Emamian, M.H., Shariati, M., Fotouhi, A., 2012. Contrast sensitivity evaluation in a population-based study in shahroud, iran. Ophthalmology 119, 541–546.

He, K., Zhang, X., Ren, S., Sun, J., 2016. Deep residual learning for image recognition, in: Proceedings of the Conference on Computer Vision and Pattern Recognition, pp. 770–778.

Hirsch, J., 1982. Falcon visual sensitivity to grating contrast. Nature 300, 57–58.

Hodos, W., Ghim, M.M., Potocki, A., Fields, J.N., Storm, T., 2002. Contrast sensitivity in pigeons: a comparison of behavioral and pattern erg methods. Documenta Ophthalmologica 104, 107–118.

Hubel, D.H., Wiesel, T.N., 1961. Integrative action in the cat’s lateral geniculate body. The Journal of Physiology 155, 385.

Hubel, D.H., Wiesel, T.N., 2004. Brain and visual perception: the story of a 25-year collaboration. Oxford University Press.

Kelly, D., 1983. Spatiotemporal variation of chromatic and achromatic contrast thresholds. JOSA 73, 742–750.

Kelly, D.H., 1979. Motion and vision. ii. stabilized spatio-temporal threshold surface. JOSA 69, 1340–1349.

Khaligh-Razavi, S.M., Kriegeskorte, N., 2014. Deep supervised, but not unsupervised, models may explain it cortical representation. PLoS computational biology 10, e1003915.

Kim, J., Lee, S., 2017. Deep learning of human visual sensitivity in image quality assessment framework, in: Proceedings of the Conference on Computer Vision and Pattern Recognition, pp. 1676–1684.

Kim, Y.J., Kim, H.S., 2010. Spatial luminance contrast sensitivity: Effects of surround. Journal of the Optical Society of Korea 14, 152–162.

Krizhevsky, A., Sutskever, I., Hinton, G.E., 2012. Imagenet classification with deep convolutional neural networks, in: Proceedings of the Conference on Advances in Neural Information Processing Systems, pp. 1097—-1105.

Kuffler, S.W., 1953. Discharge patterns and functional organization of mammalian retina. Journal of neurophysiology 16, 37–68.

Li, Q., Gomez-Villa, A., Bertalmío, M., Malo, J., 2022. Contrast sensitivity functions in autoencoders. Journal of Vision 22, 8–8.

Lin, T.Y., Maire, M., Belongie, S., Hays, J., Perona, P., Ramanan, D., Dolĺar, P., Zitnick, C.L., 2014. Microsoft coco: Common objects in context, in: European Conference on Computer Vision, Springer. pp. 740–755.

Lindsay, G., 2020. Convolutional neural networks as a model of the visual system: Past, present, and future. Journal of Cognitive Neuroscience 33, 1–15.

Marr, D., 1982. Vision: A Computational Approach. MIT press.

Mullen, K.T., 1985. The contrast sensitivity of human colour vision to redgreen and blue-yellow chromatic gratings. The Journal of physiology 359, 381–400.

Müllner, D., 2013. fastcluster: Fast hierarchical, agglomerative clustering routines for r and python. Journal of Statistical Software 53, 1–18.

Neri, P., 2022. Deep networks may capture biological behavior for shallow, but not deep, empirical characterizations. Neural Networks 152, 244–266.

Northmore, D., Dvorak, C., 1979. Contrast sensitivity and acuity of the goldfish. Vision research 19, 255–261.

Olshausen, B.A., Field, D.J., 1996. Emergence of simple-cell receptive field properties by learning a sparse code for natural images. Nature 381, 607– 609.

Owsley, C., 2003. Contrast sensitivity. Ophthalmology Clinics of North America 16, 171–177.

Peli, E., 1990. Contrast in complex images. JOSA A 7, 2032–2040.

Peli, E., Arend, L., Labianca, A.T., 1996. Contrast perception across changes in luminance and spatial frequency. JOSA A 13, 1953–1959.

Pelli, D.G., Bex, P., 2013. Measuring contrast sensitivity. Vision research 90, 10–14.

Radford, A., Kim, J.W., Hallacy, C., Ramesh, A., Goh, G., Agarwal, S., Sastry, G., Askell, A., Mishkin, P., Clark, J., et al., 2021. Learning transferable visual models from natural language supervision, in: International Conference on Machine Learning, pp. 8748–8763.

Ramanath, R., Snyder, W.E., 2003. Adaptive demosaicking. Journal of Electronic Imaging 12, 633–642.

Reymond, L., Wolfe, J., 1981. Behavioural determination of the contrast sensitivity function of the eagle aquila audax. Vision Research 21, 263– 271.

Schade, O.H., 1956. Optical and photoelectric analog of the eye. JOSA 46, 721–739.

Shelhamer, E., Long, J., Darrell, T., 2017. Fully convolutional networks for semantic segmentation. IEEE transactions on pattern analysis and machine intelligence 39, 640–651.

Srivastava, S., Wang, W., Eckstein, M., 2022. A feedforward network with a few million neurons learns from images to covertly attend to contextual cues like human and bayesian ideal observers. articlePsyArXiv preprint.

Storrs, K.R., Kietzmann, T.C., Walther, A., Mehrer, J., Kriegeskorte, N., 2021. Diverse deep neural networks all predict human inferior temporal cortex well, after training and fitting. Journal of Cognitive Neuroscience 33, 2044–2064.

Stromeyer, C., Kranda, K., Sternheim, C., 1978. Selective chromatic adaptation at different spatial frequencies. Vision research 18, 427–437.

Tang, S., Lee, T.S., Li, M., Zhang, Y., Xu, Y., Liu, F., Teo, B., Jiang, H., 2018. Complex pattern selectivity in macaque primary visual cortex revealed by large-scale two-photon imaging. Current Biology 28, 38–48.

Thomson, A.M., 2010. Neocortical layer 6, a review. Frontiers in neuroanatomy 4, 13–13.

Uhlrich, D.J., Essock, E.A., Lehmkuhle, S., 1981. Cross-species correspondence of spatial contrast sensitivity functions. Behavioural Brain Research 2, 291–299.

Vaswani, A., Shazeer, N., Parmar, N., Uszkoreit, J., Jones, L., Gomez, A.N., Kaiser, L-., Polosukhin, I., 2017. Attention is all you need, in: Proceedings of the Conference on Advances in Neural Information Processing Systems, pp. 6000—-6010.

de Vries, J.P., Akbarinia, A., Flachot, A., Gegenfurtner, K.R., 2022. Emergent color categorization in a neural network trained for object recognition. Elife 11, e76472.

Wandell, B.A., 1995. Foundations of vision. Sinauer Associates.

Wang, Z., Bovik, A.C., Sheikh, H.R., Simoncelli, E.P., 2004. Image quality assessment: from error visibility to structural similarity. IEEE transactions on image processing 13, 600–612.

Williams, D.R., Collier, R., 1983. Consequences of spatial sampling by a human photoreceptor mosaic. Science 221, 385–387.

Wuerger, S., Ashraf, M., Kim, M., Martinovic, J., Pérez-Ortiz, M., Mantiuk, R.K., 2020. Spatio-chromatic contrast sensitivity under mesopic and photopic light levels. Journal of Vision 20, 23–23.

Yamins, D.L., Hong, H., Cadieu, C.F., Solomon, E.A., Seibert, D., DiCarlo, J.J., 2014. Performance-optimized hierarchical models predict neural responses in higher visual cortex. Proceedings of the national academy of sciences 111, 8619–8624.

Yue, X., Pourladian, I.S., Tootell, R.B., Ungerleider, L.G., 2014. Curvature-processing network in macaque visual cortex. Proceedings of the National Academy of Sciences 111, E3467–E3475.

Zamir, A.R., Sax, A., Shen, W., Guibas, L.J., Malik, J., Savarese, S., 2018. Taskonomy: Disentangling task transfer learning, in: Proceedings of the Conference on Computer Vision and Pattern Recognition, pp. 3712–3722.

Zeman, A.A., Ritchie, J.B., Bracci, S., Op de Beeck, H., 2020. Orthogonal representations of object shape and category in deep convolutional neural networks and human visual cortex. Scientific reports 10, 1–12.

